# Pregnenolone and AEF0117 block cannabinoid-induced hyperlocomotion through GSK3β signaling at striatopallidal neurons

**DOI:** 10.1101/2025.03.07.641990

**Authors:** G. Tomaselli, L. Bellocchio, PL. Raux, V. Roullot-Lacarrière, A. Busquets-Garcia, V. Lalanne, Y. Mariani, A. Cathala, M. Zanese, A. Cannich, A. Grel, D. Gisquet, M. Mondesir, M. Metna-Laurent, G. Drutel, G. Marsicano, PV. Piazza, JM. Revest, M. Vallée

## Abstract

Administration of Δ^9^-tetrahydrocannabinol (THC), the main psychoactive component of the plant *Cannabis sativa*, can induce psychotic symptomatology in humans and a large spectrum of acute psychotic-like behaviors in mice, including hyperlocomotion observed at low dose of THC (0.3 mg/kg). The cellular and molecular substrates of this effect have not been fully identified yet. Here we demonstrate that THC-induced hyperlocomotion depends on plasma membrane CB1R, which regulate the β-arrestin 1/Akt/GSK3β signaling pathway in D2R-positive neurons of the dorsal striatum forming the striatopallidal pathway of the basal ganglia. Pregnenolone (PREG) and its clinically developed analog, AEF0117, which are signaling specific inhibitors of CB1R (CB1-SSi), prevented GSK3β-dependent psychomotor stimulation induced by THC. Overall, this work highlights a novel intracellular mechanism of CB1R, thereby revealing a neuronal pathway underlying an important but still underexplored effect of THC and cannabis consumption, which could help the development of innovative therapeutic concepts against psychotic conditions.

## Introduction

*Cannabis sativa* is one of the most commonly used drugs of abuse in Europe and worldwide ^1,2^, and its consumption represents a high-risk factor for the development of cannabis use disorders (CUD) ^3^. As part of CUD, converging epidemiological and clinical evidence supports an association between cannabis use and the risk of developing psychosis and schizophrenia-like diseases in humans ^4–9^. For example, the acute administration of Δ^9^-tetrahydrocannabinol (THC), which is the main psychoactive component of cannabis, can induce transient cannabis-associated psychotic-like symptoms ^8,10–12^. Accordingly, acute THC administration dose-dependently induces a spectrum of behavioral phenotypes that may recapitulate some addictive and toxic effects of cannabis use in experimental animals ^13^, including THC-induced acute psychotic-like states (CIAPS) ^14^. THC exerts dose-dependent biphasic behavioral effects on locomotor activity, food intake, cognitive function, and reward processes ^14–17^. In particular, whereas high doses decrease locomotion, low doses of THC (< 0.5mg/kg) can produce a stimulation of locomotor activity in rodent models ^14,17,18^. Although type-1 cannabinoid receptors (CB1R) have long been identified as key mediators of the psychoactive effects of THC in the brain, including locomotor hyperactivity ^17,19^, the specific downstream cellular processes have not been fully elucidated. Given the widespread prevalence of cannabis use, it appears more and more important to have a full understanding of the cellular, molecular and signaling mechanisms involved in the acute psychotropic effects of cannabis.

CB1R are among the most abundant members of the seven transmembrane G-protein-coupled receptor (GPCR) family in the mammalian brain. Upon activation by exogenous (*e.g.* THC) or endogenous agonists (so-called endocannabinoids), CB1R activate a variety of intracellular signaling cascades that are mediated by G-proteins but also by β-arrestins. A hyperactivity of the CB1R-mediated signaling has been involved in several pathological states involving both the brain and peripheral organs. Despite the large therapeutical potential of an inhibition of the CB1R, orthosteric antagonists of this receptor induce behavioral side effects which forbid their use as therapeutic tools in humans ^20–22^. The discovery of a biased inhibition of the CB1R, naturally used by the brain, through the neurosteroid pregnenolone (PREG) to control the hyperactivity of this receptor has opened a new avenue of research for the development of better tolerated CB1 inhibitors ^13^, including CIAPS in mice ^14^. Indeed, the stable and well absorbed PREG derivative AEF0117 has been shown to lack the behavioral side effects of CB1 antagonists in both animals and humans, but to block most of the behavioral effects of THC, including the CIAPS in mice, non-human primates and humans ^23^. AEF0117 and PREG have been defined as signaling specific inhibitors of the CB1R (CB1-SSi) because they do not modify agonist binding but block cannabinoid-induced activation of the ERK1/2^MAPK^ (extracellular signal-regulated kinase/mitogen-activated protein kinase) pathway, without modifying G-protein-mediated signaling such as the CB1R-dependent reduction of cyclic adenosine monophosphate (cAMP) ^13,23^. Biased inhibition of the CB1R by CB1-SSi introduces the idea that the multiple signaling pathways activated by the GPCR should not be viewed as an integrated signal, but that each molecular pathway may have different physiological roles that can be targeted independently. As a consequence, understanding the physiological role of the precise intracellular signaling mediated by GPCR becomes an important new avenue of research. Unfortunately, knowledge in this domain is largely lacking.

Here, combining complementary *in vitro* / *in vivo* pharmacological, viral, and intracellular signaling approaches, together with genetic mouse models, we investigated CB1-dependent signaling pathways and the cellular localization that mediate the locomotor hyperactivity induced by a low dose of THC. The results show that the CB1R-mediated activation of the β-arrestin 1/Akt/GSK3β signaling pathway in D2R-positive neurons of the dorsal striatum is necessary for THC-induced hyperlocomotion.

## Results

### THC-induced hyperlocomotion is specifically associated to CB1-mediated Akt-GSK3β signaling activity in the dorsal striatum of mice

We have previously shown that the biased inhibition of the CB1R by PREG or AEF0117 is able to inhibit the hyperactivity induced by low doses of THC ^14,23^. Since these drugs do not modify the main G-protein-mediated signaling of the CB1R, we focused on signaling pathways that can be modified by the CB1R in a G-protein-independent manner (Fig. 1a) ^19,24–27^. These pathways were studied in the CB1R dorsal striatum, which is a critical brain structure for motor control and psychotic-related disorders ^28,29^.

**Fig. 1.**
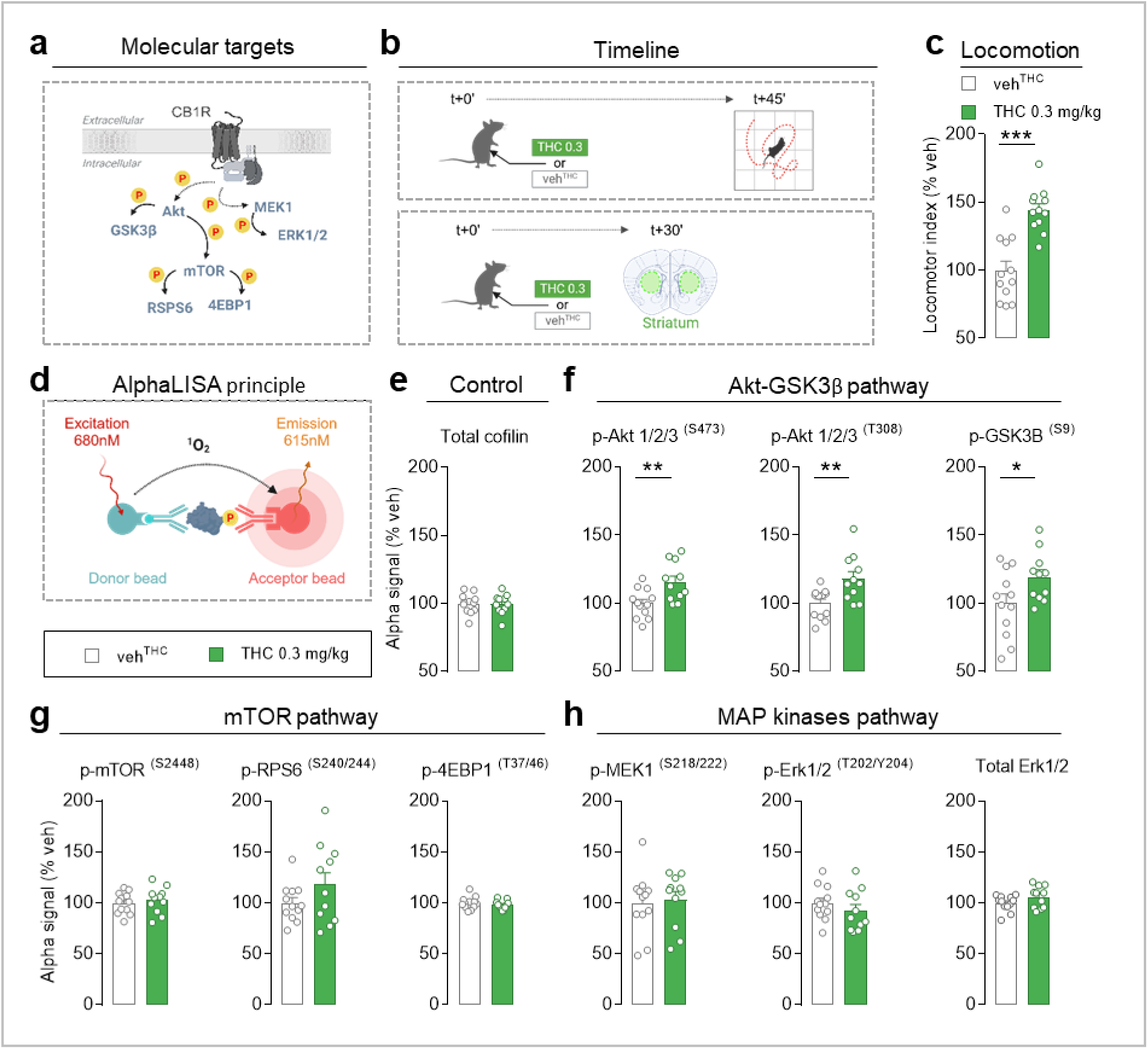
THC-induced hyperlocomotion is specifically associated to CB1R-mediated Akt-GSK3β signaling activity in the dorsal striatum of mice. **a.** Scheme of intracellular signaling pathways mediated by CB1R through sequential phosphorylation of kinases, including Akt and GSK3β and Akt-downstream kinases mTOR, RPS6, and 4EBP1, as well as MEK1 and Erk1/2 kinases. CB1R, type-1 cannabinoid receptor; Akt, protein kinase B; GSK3β, glycogen synthase kinase 3 beta; mTOR, mammalian target of rapamycin; RPS6, 40S ribosomal protein S6; 4EBP1, eukaryotic translation factor 4E-binding protein 1; MEK1, mitogen-activated protein kinase 1, extracellular signal-regulated protein kinase 1/2 (ERK1/2). **b.** Timeline of locomotor activity assayed in an open-field (top) and dorsal striatum sampling (bottom) 45 min and 30 min after i.p. injection of THC (0.3 mg/kg) or vehicle (veh^THC^), respectively. **c.** Effect of THC on locomotion. A locomotor index including horizontal and vertical locomotion (crosses + rearings) expressed as a percentage of the veh^THC^ was calculated. Two-tailed unpaired t-test; t=5.46, df=22, ***p<0.001 compared to veh^THC^. **d.** Scheme of the principle of AlphaLISA technology for the quantitative detection of phosphorylated proteins. A detailed description of the principle has been reported previously (see text). **e-h.** AlphaLISA assays of total and phosphorylated proteins from dorsal striatum protein extracts of mice treated with veh^THC^ or THC. Alpha signal is expressed as a percentage of veh^THC^. **e.** AlphaLISA assays of total cofilin used as a loading control. **f.** AlphaLISA assays of phosphorylated Akt at Ser473 and Thr308 and of GSK3β at Ser9. Two-tailed unpaired t-test; THC vs. veh^THC^, p-Akt^T308^: t=3.13, df=21, **p<0.01; p-Akt^S473^: t=3.04, df=21, **p<0.01; p-GSK3β^S9^: t=2.14, df=21, *p<0.05 compared to veh^THC^. **g.** AlphaLISA assays of phosphorylated mTOR at Ser2448, RP-S6 at Ser240/244 and of 4EBP1 at Thr37/46. Two-tailed unpaired t-test; THC vs. veh^THC^, p-mTOR^S2448^: t=0.53, df=21, p=0.6; p-RPS6^S240/244^: t=1.44, df=21, p=0.16; 4EBP1^T37/46^: t=0.88, df=21, p=0.39. **h.** AlphaLISA assays of phosphorylated MEK1 at Ser218/222 and Erk1/2 at Thr202 and Tyr204, and the total Erk1/2 protein. Two-tailed unpaired t-test; THC vs. veh^THC^, p-MEK1^S218/222^: t=0.86, df=21, p=0.39; p-Erk1/2T^202/Y204^: t=0.97, df=21, p=0.34; total Erk1/2: t=1.54, df=21, p=0.13. Data are mean ± SEM of single values.

Mice were injected intraperitoneally (i.p.) with a low dose of THC (0.3 mg/kg), known to activate locomotor activity in a novel environment ^14^ (Fig. 1b). Forty-five minutes after injection, we observed THC-induced hyperlocomotion in an open-field maze compared to vehicle-treated mice (Fig. 1c). This behavioral hyperactivity was spatio-temporally consistent with increased plasma and striatal THC levels in mice following i.p. THC administration (Fig. S1).

In order to identify intracellular signaling pathways mediating THC-induced hyperlocomotion, a new batch of mice was sacrificed 30 minutes after THC injection and their dorsal striatum was extracted (Fig. 1b), and some of the major signaling pathways modulated by CB1R ligands were analyzed using a validated high-throughput technique based on the sensitive and reproducible state-of-the-art Alpha (Amplified Luminescent Proximity Homogeneous Assay) approach. This method provides the detection and measurement of the phosphorylation levels of specific proteins in small areas of the mouse brain, allowing the analysis of a large number of signaling pathways in the same brain sample ^30^ (Fig. 1d). The housekeeping protein cofilin was chosen as a loading control to ensure data reliability in all Alpha assays, and to assess whether differences between phosphorylated targets were solely the result of THC treatment ^30^. Indeed, we found that cofilin levels were not altered by THC injection (Fig. 1e). We found that administration of THC (0.3 mg/kg) selectively increased the phosphorylation of the serine/threonine kinase Akt at two phosphorylation sites, the amino residues serine 473 (Akt^S473^) and threonine 308 (Akt^T308^) (Fig. 1f). THC (0.3 mg/kg) also increased the phosphorylation of Akt downstream kinase GSK3β by phosphorylating the serine 9 residue (GSK3β^S9^) (Fig. 1f), an effect that results in the inhibition of the activity of GSK3β ^31^. In contrast, there was no change in the mTOR cascade, which can also be initiated by Akt (Fig. 1a). THC did not induce the phosphorylation of mTOR at serine 2448 (mTOR^S2448^), nor the phosphorylation of mTOR downstream effectors RPS6 (40S ribosomal protein S6) at serine residues 240 and 244, and 4EBP1 (eukaryotic translation factor 4E-binding protein 1) at threonine residues 37 and 46 (Fig. 1g). Finally, THC did not modify the phosphorylation of Erk1/2^MAPK^ kinase, a prototypical marker of the MAPK pathway activation in mammalian cells, and the upstream MEK1 (MAP/ERK kinase-1) protein (Fig. 1a; Fig. 1h), a serine-threonine/tyrosine kinase phosphorylated at serine residues 218 and 222 (MEK1^S218/222^), which is responsible for the activation of ERK1/2^MAPK^ at both threonine 202 (Erk1/2^T202^) and tyrosine 204 (Erk1/2^Y204^) phosphorylation sites ^32^. These data clearly show that the i.p. injection of 0.3 mg/kg THC selectively engages the Akt-GSK3β pathway without modifying other important CB1R-mediated phosphorylation pathways, suggesting that THC-induced hyperlocomotion is associated with activation of Akt and subsequent inhibition of GSk3β signaling in the dorsal striatum of mice.

In a second series of experiments, we investigated whether an alteration of the Akt-GSK3β pathway may play a causal role in THC-induced hyperlocomotion. The brain-penetrating drug GDC-0084 is an inhibitor of PI3-kinase (PI3K), which is an upstream target of Akt-GSK3β signaling ^33^. GDC-0084 was shown to decrease p-Akt^S473^ in mouse brain tissue 1-hour after administration ^34^. First, we determined the highest sub-effective dose of GDC-0084 that did not alter locomotion *per se. Per os* (p.o.) administration of GDC-0084 decreased locomotion in a dose-dependent manner, and 6 mg/kg resulted the highest ineffective dose on locomotor activity (Fig. S2). In order to determine whether the Akt-GSK3β signaling mediates the hyperlocomotor effect of THC, 6 mg/kg GDC-0084 was administered 1 hour before the cannabinoid treatment and locomotion or molecular signaling in striatal cells was assessed 45 min and 30 min later in separate groups of mice (Fig. 2a). We found that GDC-0084 selectively prevented both THC-induced hyperlocomotion (Fig. 2b) and THC-induced increases in phosphorylation/activation of Akt at T308 and S473 and phosphorylation/inhibition of GSK3β at S9 in the dorsal striatum (Fig. 2c), without affecting locomotion (Fig. S2 and Fig. 2b) or GSK3β^S9^ phosphorylation *per se* (Fig. 2c). Although GDC-0084 has also been described as an mTOR inhibitor ^33^, this inhibition is unlikely to underlie the rescue of THC-behavioral effects by GDC-0084, as we did not find any changes in the mTOR signaling pathway in the dorsal striatum of THC-treated mice (Fig. 1g). Taken together, our data demonstrate that CB1R-mediated activation of GSK3β signaling is a necessary condition to induce hyperlocomotor activity in response to a low dose of THC in mice.

**Fig. 2.**
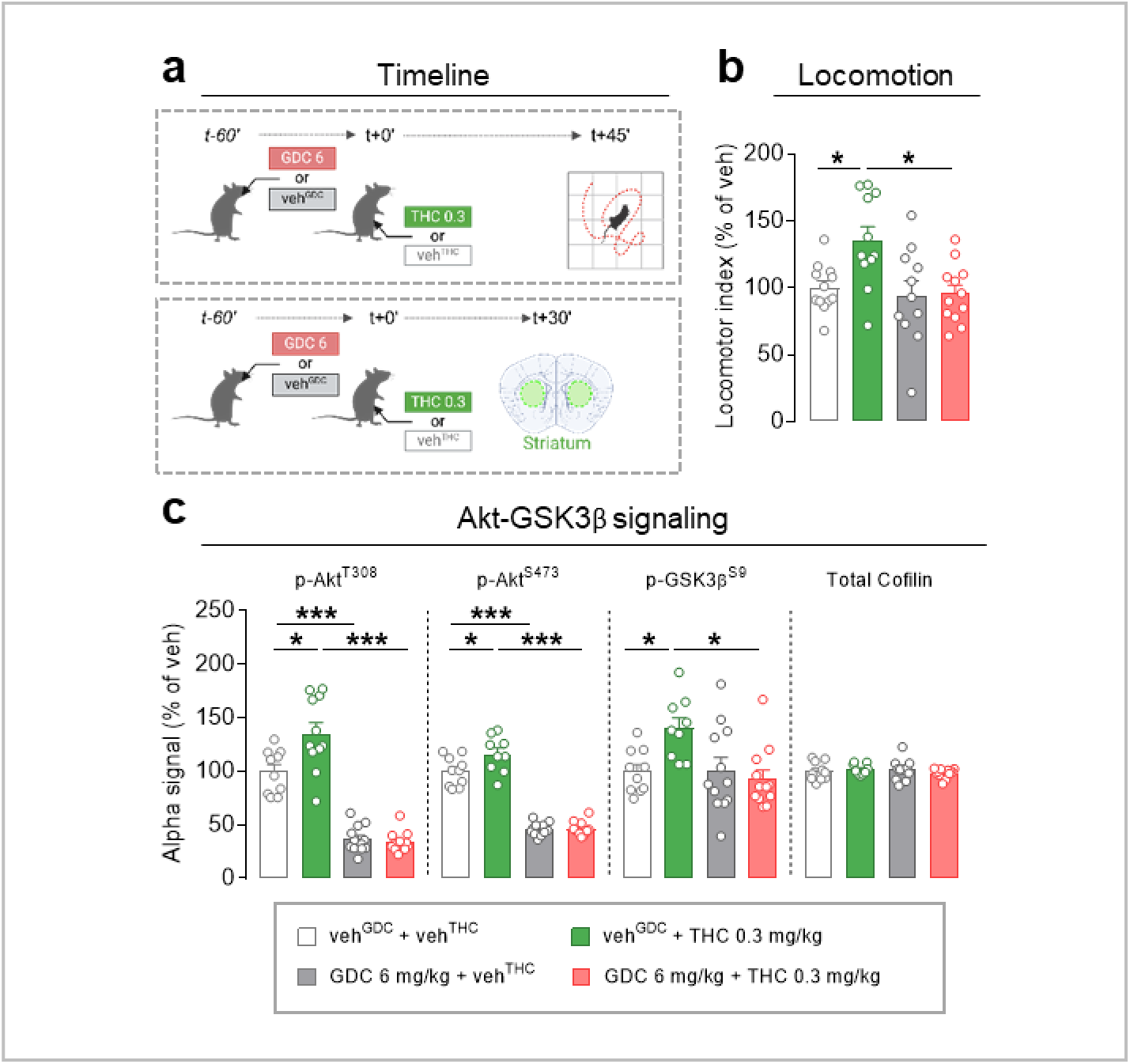
GDC-0084 blocks the effects of THC on locomotion and Akt-GSK3β pathway activity in mice. **a**. Timeline of locomotor activity assayed in an open-field (top) and dorsal striatum sampling (bottom) 45 min and 30 min after i.p. injection of THC (0.3 mg/kg) or vehicle (veh^THC^), respectively with a 1 h pre-treatment of oral administration of GDC-0084 (6 mg/kg) or vehicle (veh^GDC^). **b**. Effect of GDC-0084 (6 mg/kg) on locomotion in mice treated with THC or veh^THC^. A locomotor index including horizontal and vertical locomotion (crosses + rearings) expressed as a percentage of the veh^THC^ was calculated. Two-way ANOVA, treatment interaction, F(1,42)=4.1, p<0.05; Tukey’s multiple comparison test, THC effect in mice treated with the vehicle of GDC-0084 (veh^GDC^), *p<0.05; GCD-0084 vs. veh^GDC^ in mice treated with veh^THC^, p=0.99 or with THC, *p<0.02. **c**. Effect of GDC-0084 (6 mg/kg) on phosphorylation of Akt at Thr308 and Ser473 and of GSK3β at Ser9, and total cofilin in dorsal striatum extracts from mice treated with THC (0.3 mg/kg) or veh^THC^. Alpha signal is expressed as a percentage of veh^THC^. Two-way ANOVA; treatment interaction, p-Akt^T308^: F(1,39)=6, p<0.02; p-Akt^S473^: F(1,39)=5.4, p<0.05; p-GSK3β^S9^: F(1,39)=6.6, p<0.02; total cofilin: F(1,39)=1.1, p=0.30; Tukey’s multiple comparison test; THC effect in mice treated with veh^GDC^, p-Akt^T308^: *p<0.05; p-Akt^S473^: *p<0.02; p-GSK3β^S9^: *p<0.05; GCD-0084 vs. veh^GDC^ in veh^THC^-treated mice, p-Akt^T308^: ***p<0.001; p-Akt^S473^: ***p<0.001; p-GSK3β^S9^: p=0.98 and in THC-treated mice, p-Akt^T308^: ***p<0.001; p-Akt^S473^: ***p<0.001; p-GSK3β^S9^: *p<0.02. Data are mean ± SEM of single values.

### β-Arrestin 1 is involved in THC-induced Akt-GSK3β signaling activity in striatal cell lines

We then further explored the CB1R-mediated intracellular cascade by analyzing which upstream signal transducer may sustain GSK3β signaling. Although the β-arrestin family, including β-arrestin 1 and 2 isoforms, has been classically implicated in the regulation of GPCR desensitization and internalization, novel functions as signal transducers are emerging. Accordingly, β-arrestin 2 has been shown to be more involved in desensitization and internalization of the receptor, while β-arrestin 1 has been involved in the initiation of signaling ^35^. We therefore investigated the role of β-arrestin 1 protein in the modulation of Akt-GSK3β signaling activity by THC. We first performed *in vitro* experiments using a cell model of murine striatal medium spiny neurons, the STH*dh*^Q7/Q7^ cell line ^36^, which is a physiologically relevant model for studying CB1R biased signaling in the striatum since these cells derive from this brain region and endogenously express CB1R. This cell line was shown to display biased CB1R signaling in response to several CB1R agonists ^37^, including β-arrestin 1-mediated phosphorylation of Akt at S473 ^38^. In contrast to what has been previously reported ^38^, the highly accurate method of ddPCR (digital droplet polymerase chain reaction) ^39^ revealed that our clone of STH*dh*^Q7/Q7^ cells contain low levels of transcripts derived from the β-Arrestin 1 encoding gene (*Arrb1*), as compared to that of other signal transducers, such as β-arrestin 2 encoding gene (*Arrb2*), and G-proteins (*Gnai1/2/3, Gnao, Gnaq, Gna11 and Gnas*) (Fig. S3a). Therefore, to induce an increase in β-Arrestin 1 protein, we transfected STH*dh*^Q7/Q7^ cells with a plasmid expressing the human β-Arrestin 1 protein (Fig. S3b). Then, STH*dh*^Q7/Q7^ cells were treated with either THC (10 µM) or vehicle solution and Akt-GSK3β signaling was assessed by AlphaLisa 30 min later (Fig. 3a). The results showed a significant increase in Akt^T308^, Akt^S473^ and GSK3β^S9^ phosphorylation in response to THC as compared to vehicle in transfected cells, while no effect of THC on signaling activity was observed in control STH*dh*^Q7/Q7^ cells transfected with the empty vector (Fig. 3b).

**Fig. 3.**
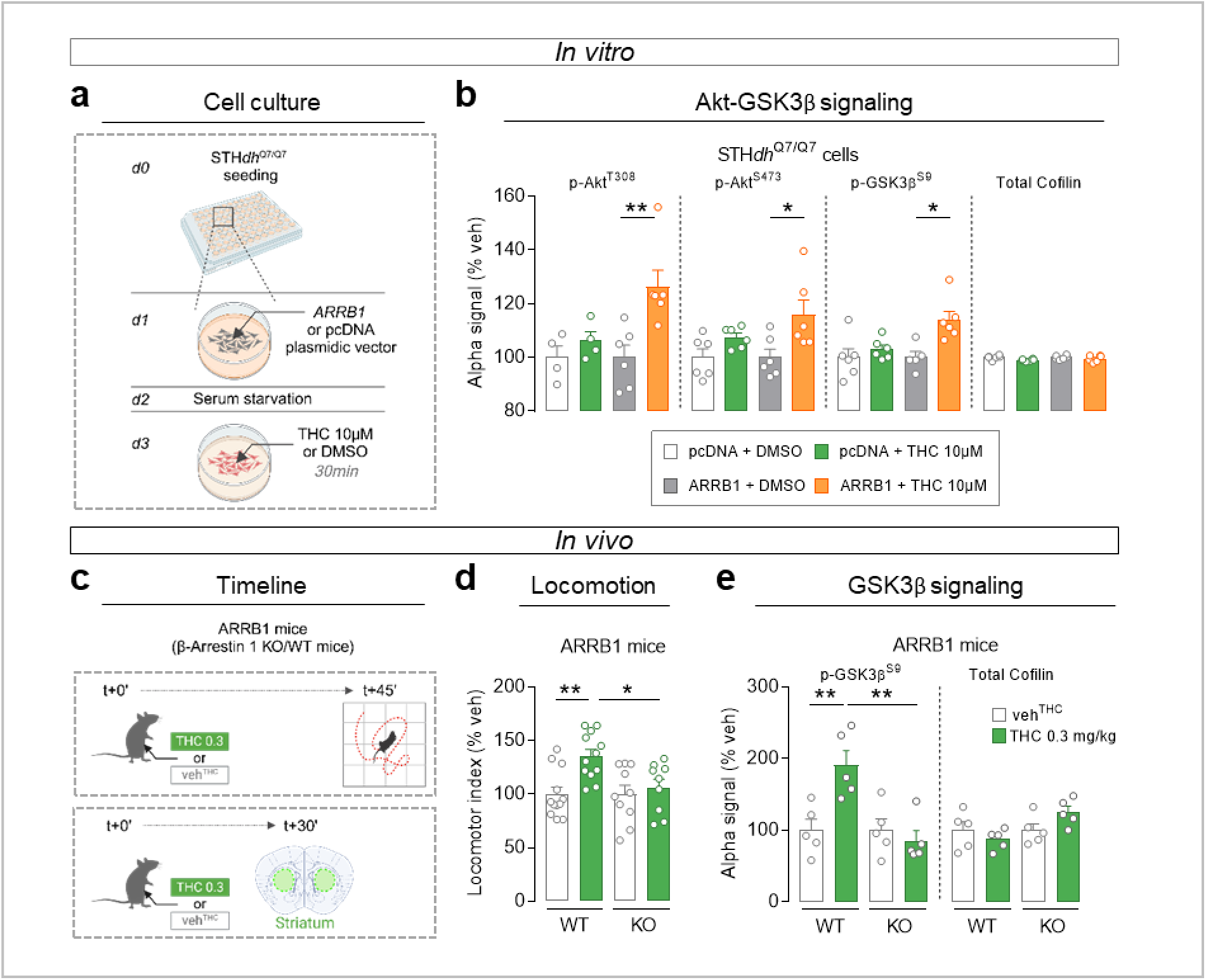
β-arrestin 1 is required for THC effects on Akt-GSK3β signaling activity in cell culture, on locomotor activity in mice, and on GSK3β phosphorylation in mouse striatal tissue. **a.** STH*dh^Q7/Q7^* cell transfection timeline. Day 0, STH*dh^Q7/Q7^* cells were seeded in 96-well plates. The day after STH*dh^Q7/Q7^* cells were transfected with *ARRB1* gene encoding human β-Arrestin 1 or empty pcDNA vectors. After 24 h of serum starvation, cells were treated with THC (10 µM) or DMSO (vehicle) for 30 min before AlphaLISA assay. **b**. Effect of THC on phosphorylation of Akt at Thr308 and Ser473, GSK3β at Ser9 and total cofilin in STH*dh^Q7/Q7^* cells transfected with pcDNA or *ARRB1* gene. Alpha signal is expressed as a percentage of vehicle. Two-way ANOVA, treatment effect, p<0.01 for p-Akt^T308^, p-Akt^S473^ and p-GSK3β^S9^, and p<0.05 for total cofilin; Tukey’s multiple comparison test; THC effect on pcDNA transfected cells, p-Akt^T308^: p=0.87, p-Akt^S473^: p=0.48; p-GSK3β^S9^: p=0.84; total cofilin: p=0.19 and on *ARRB1* transfected cells, p-Akt^T308^: **p<0.01; p-Akt^S473^: *p<0.05; p-GSK3β^S9^: *p<0.02; total cofilin: p=0.78. **c**. Timeline of locomotor activity assayed in an open-field (top) and dorsal striatum sampling (bottom) 45 min and 30 min after i.p. injection of THC (0.3 mg/kg) or vehicle (veh^THC^) in β-arrestin 1 knock-out (KO; Arrb1 ^-/-^) mice and wild-type (WT) littermates. **d**. Effect of THC on locomotor activity in WT and KO mice. A locomotor index including horizontal and vertical locomotion (crosses + rearings) expressed as a percentage of veh^THC^ was calculated. Two-way ANOVA, genotype x treatment interaction, F(1,39)=4.2, p<0.05; Tukey’s multiple comparison test; THC vs. veh^THC^ in WT and KO mice: **p<0.01 and p=0.93, respectively; THC effect in WT vs KO mice: *p<0.05. **e**. Effect of THC on phosphorylation of GSK3β at Ser9 and total cofilin in dorsal striatum extracts from WT and KO mice. Alpha signal is expressed as a percentage of veh^THC^. Two-way ANOVA, genotype x treatment effect, p-GSK3β^S9^: F(1,16)=10, p<0.01, total cofilin: F(1,16)=4.3, p=0.054; Tukey’s multiple comparison test for p-GSK3β^S9^; THC vs. veh^THC^ in WT and KO mice, **p<0.01 and p>0.92, respectively; THC effect in WT vs KO mice: **p<0.01. Data are mean ± SEM of single values.

### β-Arrestin 1 mediates THC-induced hyperlocomotion and increased GSK3β phosphorylation in mice

To further confirm *in vivo* that β-Arrestin 1 triggers the Akt-GSK3β pathway to induce the hyperlocomotor effect of THC, we performed behavioral experiments using β-Arrestin 1-deficient (Arrb1^-/-^) mice ^40^. Locomotion was assessed on adult male Arrb1^-/-^ mice and wild-type (WT) littermates 45 min after THC injection at a dose of 0.3 mg/kg (Fig. 3c). Strikingly, whereas locomotor activity was increased in THC-treated WT mice, the cannabinoid drug had no effect in Arrb1^-/-^ mice (Fig. 3d). The lack of increased locomotion in Arrb1^-/-^ mice was specific to THC, as amphetamine-induced hyperlocomotion was maintained in both genotypes (Fig. S4), thereby confirming that β-Arrestin 1 is indeed necessary to mediate THC-induced hyperlocomotion. Finally, we found an increase in phosphorylated GSK3β^S9^ in response to THC in the dorsal striatum of WT mice, but not in Arrb1^-/-^ littermates (Fig. 3e), further confirming the causal role of β-Arrestin 1 in the modulation of the Akt-GSK3β pathway by THC.

### THC-induced hyperlocomotion involves CB1R located on striatal D2 neurons

Next, we investigate the neuronal networks involved in THC-induced hyperlocomotion in the dorsal striatum. Besides the classical localization at the plasma membrane typical of all GPCRs, CB1R are also intracellularly associated with different organelles, including mitochondria (mtCB1R), where they control several cellular functions, such as respiration, ATP production and calcium homeostasis ^41,42^. Importantly, it has recently been shown that striatal mtCB1R are responsible for the hypolocomotor effect of high doses of CB1R agonists ^45^. Therefore, we asked whether mtCB1R are involved in the hyperlocomotor effect of low doses of THC. Using the newly generated and characterized mutant mouse line lacking mtCB1R, the DN22-CB1-KI mice ^45^, we showed a significant and similar THC effect on locomotion in mutant and WT littermates as compared to vehicle (Fig. 4a), suggesting that mtCB1R is not involved in the increased locomotor activity. To dissect the cellular location of the CB1R involved in THC-induced hyperlocomotion, we then focused on CB1R expressed in striatal medium spiny neurons (MSN) ^44–46^. Abundant pools of CB1R are most commonly found presynaptically at the terminals of MSNs projecting to the globus pallidus or to substantia nigra reticulata ^47–49^. Using a conditional mutant mice lacking CB1R in GABAergic neurons, including MSNs (hereafter named GABA-CB1-KO mice) ^15,45^, we found indeed that THC was unable to induce hyperlocomotion in the GABA-CB1-KO mice, when compared to their WT (CB1-flox) littermates (Fig. 4b). This suggests that CB1R expressed in GABA neurons, including MSNs, are necessary for the enhanced locomotion induced by THC. The two subtypes of MSNs, designated as direct (striatonigral) and indirect (striatopallidal) pathways, have been extensively described to play a role in the control of movement and action ^50–53^. The direct pathway originates from MSNs expressing the dopamine 1 receptor (D1-MSNs) projecting primarily to the substantia nigra reticulata (SNr) and globus pallidus internae (GPi), and the indirect pathway originates from MSNs expressing dopamine D2 (D2-MSNs) and adenosine A2a receptors projecting to the globus pallidus externae (GPe), which in turn can target the subthalamic nucleus (STN) and GPi/SNr ^50,54^. CB1R subpopulations may be involved in the functioning of the direct and indirect pathways and in the control of various aspects of motor behavior ^53^. Therefore, we tested whether CB1R expressed by D1-MSNs and/or D2-MSNs may underlie the hyperlocomotor effect of THC. To this aim, we used the validated conditional mouse lines Drd1-CB1-KO and Adora2a-CB1-KO, which lack the CB1R in D1-MSNs or in D2-MSNs, respectively ^46^. THC-induced hyperlocomotion was still present in Drd1-CB1-KO mice as in their WT littermates (Fig. 4c), but it was abolished in Adora2a-CB1-KO mice (Fig. 4d). This result strongly indicates that CB1R at D2-MSNs rather than D1-MSNs is necessary for enhanced locomotion in response to low doses of THC. The lack of hyperactivity in Adora2a-CB1-KO mice was not due to a motor deficit as it was previously shown that Adora2a-CB1-KO mice exhibit normal motor activity, coordination, and amphetamine-induced hyperlocomotion ^46^. Since specific reduction in CB1 protein levels was shown in the GPe of Adora2a-CB1-KO mice compared to WT littermates ^46^, our data suggest a presynaptic inhibition by CB1R at D2-MSNs as a possible neurophysiological mechanism for THC-induced hyperlocomotion. According to this hypothesis, potentiation of D2-MSNs activity by the D2R antagonist haloperidol ^29^ was able to counteract the increase in locomotor activity induced by THC (Fig. S5). We further explored GSK3β-phosphorylation at S9 by Akt in D2-MSNs as a possible cellular and molecular substrate for THC-induced hyperlocomotion. To this aim, we injected a viral vector (*i.e.,* AAV-DIO-GSK3β^S9A^) expressing a Cre-dependent HA-tagged dominant negative mutant form of GSK3β bearing a point mutation that converts serine 9 to alanine (*i.e.,* GSK3β^S9A^) thus preventing its phosphorylation, in the dorsal striatum of Tg(Adora2a*cre) KG139Gsat mouse line (Fig. 4e). We first confirmed, by HA-immunodetection coupled to fluorescent *in situ* hybridization against D1R or D2R, that the dominant negative GSK3β^S9A^ protein (in green) was not expressed in D1 expressing neurons (labeled in red in left panel) whereas it was selectively expressed in D2-expressing neurons as shown by the yellow colocalization (right panel) (Fig. 4f). Mice were then injected with THC (0.3 mg/kg) and locomotor activity was recorded 45 minutes afterwards. We found that virally disabling GSK3β phosphorylation (*i.e.,* GSK3β^S9A^) in D2-MSNs prevented THC-induced hyperlocomotion (Fig. 4g). Overall, our findings demonstrated that GSK3β inactivation is a sufficient condition to mediate THC-induced hyperlocomotion through CB1R at D2-MSNs.

**Fig. 4.**
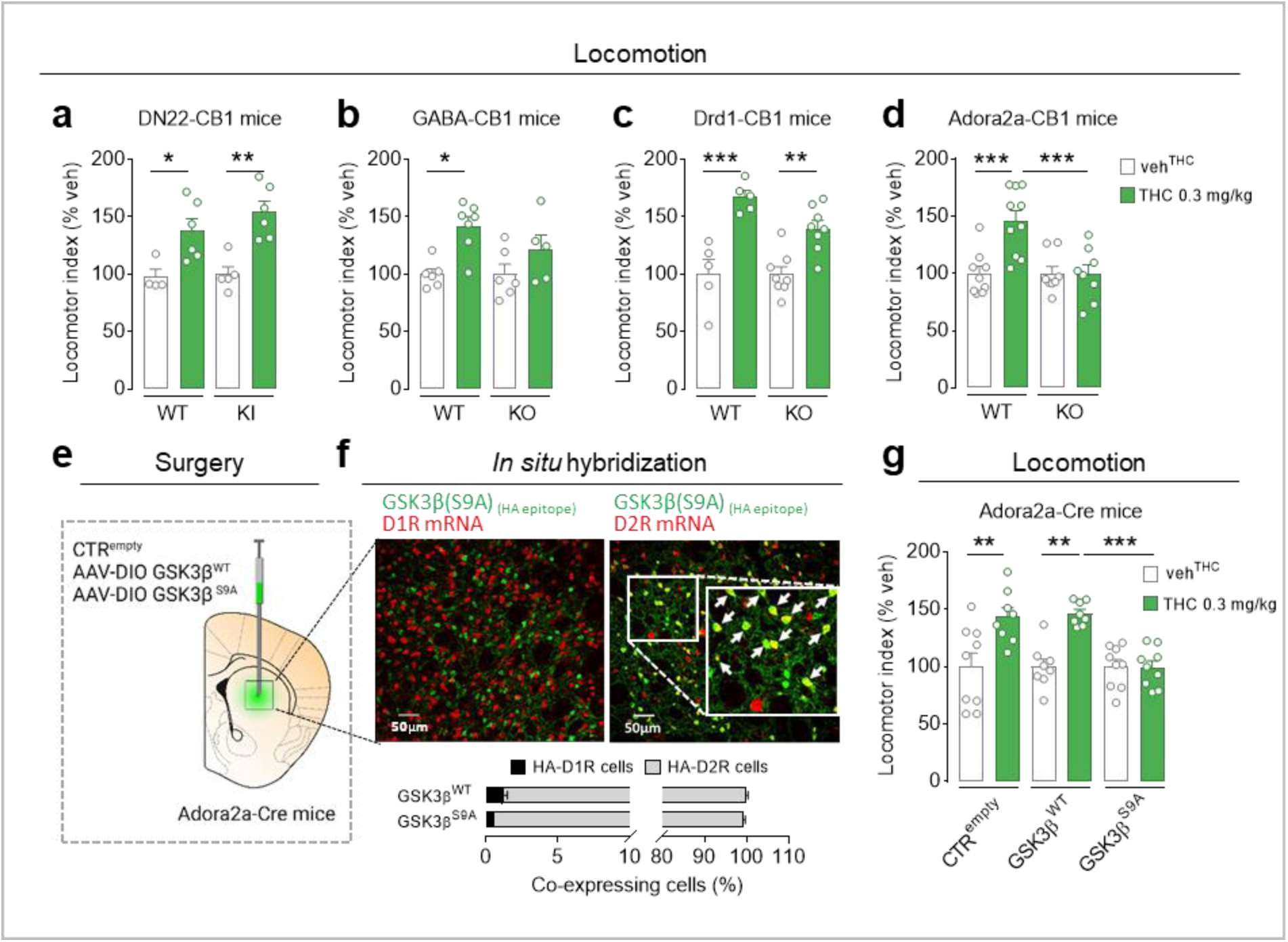
CB1R located on plasma membrane of striatal D2 neurons trigger THC-induced hyperlocomotion in mice. **a-d, g.** Effect of THC on locomotor activity in mutant mice and wild-type (WT) littermates. The locomotor index, including horizontal and vertical locomotion (crosses + rearings), is expressed as a percentage of the vehicle (veh^THC^). **a**. Locomotion in DN22-CB1-KI mice, lacking mitochondrial CB1R (mtCB1R). Two-way ANOVA, genotype effect, F(1,17)=1, p=0.32; treatment effect, (F(1,17)=27, p<0.001; Tukey’s multiple comparison test, THC vs. veh^THC^ in WT and DN22-CB1-KI mice, *p<0.05 and **p<0.01, respectively. **b**. Locomotion in GABA-CB1-KO mice, lacking CB1R on GABAergic neurons. Two-way ANOVA, genotype effect, F(1,20)=1.4, p=0.25; treatment effect, F(1,20)=13, p<0.01; Tukey’s multiple comparison test, THC vs. veh^THC^ in WT and GABA-CB1-KO mice, *p<0.01 and p=0.36, respectively. **c**. Locomotion in Drd1-CB1-KO mice, lacking the CB1R at the terminals of D1-MSNs (in the substantia nigra). Two-way ANOVA, genotype effect, F(1,22)=2.8, p=0.10; treatment effect, (F(1,22)=, p<0.001; Tukey’s multiple comparison test, THC vs. veh^THC^ in WT and Drd1-CB1-KO mice, ***p<0.001 and **p<0.01, respectively. **d**. Locomotion in Adora2a-CB1-KO mice, lacking the CB1R at the terminals of D2-MSNs (in the globus pallidus externae). Two-way ANOVA, interaction treatment x genotype, F(1,31)=9.2, p<0.01; Tukey’s multiple comparison test, THC vs. veh^THC^ in WT and Adora2a-CB1-KO mice, ***p<0.001 and p>0.99, respectively; THC effect in WT vs Adora2a-CB1-KO mice, ***p<0.001. **e**. Surgical scheme for viral injection of control virus (CTR^empty^), AAV-DIO-GSK3β^WT^ or AAV-DIO-GSK3β^S9A^ in the dorsal striatum of Adora2a-Cre mice. **f**. Illustrations of immunofluorescence and double fluorescent *in situ* hybridization experiments (top). GSK3β^S9A^ (green) and D1R mRNA (left) and D2RmRNA (right) (red). Scale bars, 50 µm. Relative quantification of co-expressing cells (yellow) of GSK3β^WT^ and GSK3β^S9A^ for HA-D1R and HA-D2R cells (bottom). **g**. Effect of THC on locomotor activity in Adora2a-Cre mice injected with CTR^empty^, GSK3β^WT^ or GSK3β^S9A^ viruses into the dorsal striatum. Two-way ANOVA, interaction treatment x viral injection, F(2,45)=6.2, p<0.01; Tukey’s multiple comparison test, THC vs. veh^THC^ in Adora2a-Cre mice injected with CTR^empty^, GSK^WT^ or GSK^S9A^ virus, **p<0.01, **p<0.01 or p>0.99, respectively. THC effect in Adora2a-Cre mice injected with GSK^WT^ vs. GSK^S9A^ virus, ***p<0.001. Data are mean ± SEM of single values.

### PREG and AEF0117 prevent THC-induced GSK3β-dependent hyperlocomotion in mice

Finally, we tested whether PREG and AEF0117 could inhibit THC-induced hyperlocomotion by acting on the CB1-dependent GSK3β pathway described here. After a 10 min subcutaneous (s.c.) pre-treatment with PREG (6 mg/kg), THC-induced hyperlocomotion and signaling were assessed 45 min and 30 min following THC injection in an open-field maze and by AlphaLisa, respectively (Fig. 5a). Confirming previous findings ^14^, the results showed that pre-treatment with PREG blocked the increase in locomotion induced by THC, but did not change basal locomotion (Fig. 5b). We then further demonstrated that PREG was able to prevent THC-induced increased GSK3β^S9^ phosphorylation, without no effect *per se* (Fig. 5c). However, PREG, in addition to being a biased CB1 inhibitor, is also the precursor of all the other steroid hormones and oral administration of PREG can induce an increase in downstream steroids in humans ^55^. Although 6 mg/kg of PREG injected subcutaneously does not seem to increase the plasma levels of other steroids in rodents ^13^, to exclude this possibility, we tested the synthetic non-metabolized PREG derivative AEF0117 under the same conditions. AEF0117, differently from PREG, is not transformed in PREG downstream steroids, has a long half-life, and is well absorbed orally and can consequently be used as a therapeutic drug ^23^. After a 180 min p.o. pre-treatment with AEF0117 (15 µg/kg), THC-induced hyperlocomotion and signaling were assessed 45 min and 30 min following THC injection in an open-field maze and by AlphaLisa, respectively (Fig. 5d). Confirming previous findings ^23^, pre-treatment with AEF0117 blocked the effect of THC (0.3 mg/kg, i.p.) on locomotion in mice, while AEF0117 did not alter locomotion *per se* (Fig. 5e). In addition, the increase in GSK3β^S9^ phosphorylation induced by THC was prevented by AEF0117, which had no effect *per se* (Fig. 5f). These data indicate that PREG and AEF0117 block the increase in locomotor activity induced by a low dose of THC and provide evidence for a novel mechanism of action of AEF0117 involving the GSK3β-dependent CB1 pathway.

**Fig. 5.**
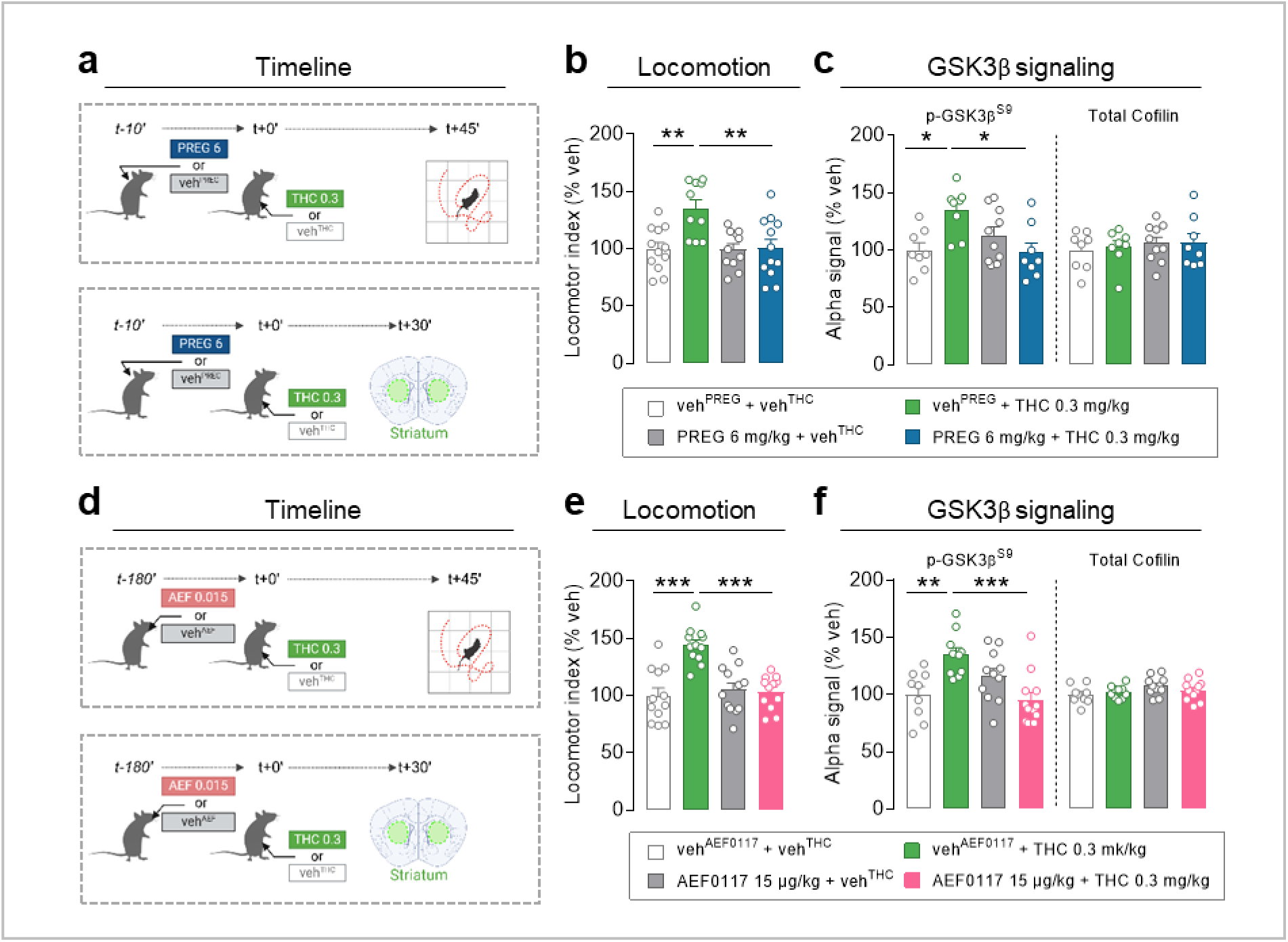
PREG and AEF0117 reverse CB1R-mediated effects of THC on locomotion and Akt-GSK3β pathway activity. **a,d.** Timeline of locomotion assayed in an open-field (top) and dorsal striatum sampling (bottom) 45 min and 30 min after i.p. injection of THC (0.3 mg/kg) or vehicle (veh^THC^), respectively, with **(a)** a 10 min pre-treatment with s.c. injection of PREG (6 mg/kg) or vehicle (veh^PREG^) or **(d)** a 3 h pre-treatment with oral administration of AEF0117 (15 µg/kg) or vehicle (veh^AEF0117^). **b,e.** A locomotor index including horizontal and vertical locomotion (crosses + rearings) was calculated. **b**. Effect of PREG on locomotion in mice treated with THC or veh^THC^. Two-way ANOVA, treatment interaction, F(1,41)=6.8, p<0.02; Tukey’s multiple comparison test, THC vs. veh^THC^ in mice treated with veh^PREG^ or PREG, **p<0.01 or p=0.99, respectively. THC effect in veh^PREG^ vs PREG-treated mice, **p<0.01. PREG effect vs. veh^PREG^ in veh^THC^-treated mice, p=0.99. **e**. Effect of AEF0117 on locomotion in mice treated with THC or veh^THC^. Two-way ANOVA, treatment interaction, F(1,44)=18, p<0.001; Tukey’s multiple comparison test, THC vs. veh^THC^ in mice treated with veh^AEF0117^ or AEF0117, ***p<0.001 and p=0.99, respectively. THC effect in veh^AEF0117^ vs AEF0117-treated mice, ***p<0.001. AEF0117 effect vs. veh^AEF0117^ in veh^THC^-treated mice, p=0.91. **c,f.** GSK3β signaling was assessed by phosphorylation of GSK3β at Ser9 (p-GSK3β^S9^). **c.** Effect of PREG on p-GSK3β^S9^ in dorsal striatum extracts from mice treated with THC or veh^THC^. Two-way ANOVA, treatment interaction, F(1,30)=11, p<0.01; Tukey’s multiple comparison test, THC vs. veh^THC^ in mice treated with veh^PREG^ or PREG, mice, *p<0.05 or p=0.60, respectively. THC effect in veh^PREG^ vs PREG-treated mice, *p<0.02. PREG effect vs. veh^PREG^ in veh^THC^-treated mice, p=0.48. **f**. Effect of AEF0117 on p-GSK3β^S9^ in dorsal striatum extracts from mice treated with THC or veh^THC^. Two-way ANOVA, treatment interaction, F(1,38)=18, p<0.001; Tukey’s multiple comparison test, THC vs. veh^THC^ in mice treated with veh^AEF0117^ or AEF0117, **p<0.01 and p=0.11, respectively. THC effect in veh^AEF0117^ vs AEF0117-treated mice, ***p<0.001. AEF0117 effect vs. veh^AEF0117^ in veh^THC^-treated mice, p=0.34. There was no difference in total cofilin between groups. Data are expressed as a percentage of veh^THC^. Data are mean ± SEM of single values.

## Discussion

In this study we show that, striatal CB1R expressed by D2-MSNs mediate the hyperlocomotor effect of a low dose of THC, through the β-arrestin 1/Akt/GSK3β signaling pathway. In addition, we demonstrated that this signaling pathway also mediate the inhibition of THC-induced hyperactivity by the neurosteroid PREG and its druggable analog, the first-in-class CB1-SSi, AEF0117.

### Relevance of THC dose

The dose of THC used in this study (0.3 mg/kg) is relevant as it reflects the psychostimulant effect of acute cannabis exposure in healthy individuals ^12^. Psychotogenic drugs cause hyperlocomotion in rodents, suggesting that this effect represents a proxy of positive symptoms of psychotic-like state ^14,56^. Indeed, the results of our behavioral experiments are consistent with those of previous studies ^14,23^ showing that THC administration at 0.3 mg/kg increased locomotor behavior in mice. In addition, plasma THC concentrations after i.p. injection reached a mean peak level of 12 ng/ml (Cmax) (Fig. S1a), which may be relevant to plasma THC levels producing impairment of neurocognitive performance, including perceptual motor control and motor inhibition, in humans ^57^. The Cmax of THC occurring at 30 min (Tmax) in mouse plasma is consistent with previous studies in rodents showing that the plasma clearance following i.p. administration is slower than with other modes of administration, such as inhalation ^58^, for which a Tmax of 10-15 min after initial administration has been reported in rodents and humans ^58,59^. As expected, Tmax was delayed in mouse striatum compared to plasma (Fig. S1b). However, we found a significant increase in THC levels at T30 min, which is relevant with the time point chosen to analyze striatal CB1 signaling components.

### Post-translational modifications, which cell signaling and where

Post-translational modifications and notably protein phosphorylation through the activities of protein kinases modifying enzymes play a central role in changing the functional properties of a protein, ultimately leading to cellular and behavioral changes ^60^. Our data demonstrate a novel CB1-mediated functional mechanism involving the phosphorylation of Akt and GSK3β-mediated β-arrestin 1 in D2-MSN neurons, which is responsible for an increase in motor behavior.

Using the high-throughput Alpha method ^30^, we unraveled a key CB1-mediated signaling pathway modulating hyperlocomotion in response to low dose of THC. Hyperlocomotion 45 min following THC administration was selectively associated with earlier (30 min post-THC) increases in Akt^T308/S473^ phosphorylation/activation and GSK3β^S9^ phosphorylation/inhibition (Fig. 1). It is worth mentioning that GSK3β^S9^ is inhibited when phosphorylated. GSK3β is a versatile “hub” kinase for GPCR-induced signaling pathways and its specificity of action can rely on the subcellular localization of GSK3β, its substrates and the distribution of its interacting receptors ^31^. Indeed, a low dose of THC inactivates CB1R-mediated GSK3β activity in striatopallidal neurons in mice by increasing its phosphorylation state. Alteration in GSK3β activity is necessary for THC-induced hyperlocomotion, as inhibition of upstream PIP3 phosphorylation/activation with GDC-0084, a potent and blood-brain barrier-penetrating PI3K inhibitor ^33,34^, specifically blocked the effect of THC on striatal phosphorylated Akt/GSK3β and on locomotor activity without having any effect *per se* (Fig. S2; Fig. 2).

Using *in vitro* and *in vivo* approaches, we demonstrated that CB1R-dependent hyperlocomotion mediated by low-dose of THC requires β-arrestin 1 as a signal transducer (Fig. 3). This is consistent with the functional selectivity of CB1R described *in vitro* involving β-arrestin 1 signaling ^38^ thereby highlighting that behavioral effects of CB1R activation can be triggered through G-protein-independent pathways to mediate a specific behavioral output ^35,41,61–63^. The study using the *arrb1* KO mice, which does not constitutively express β-arrestin 1, further confirms that CB1R-mediated GSK3β-dependent psychomotor stimulation requires a β-arrestin 1-biased modulation.

The dose-dependent effects of THC, including locomotor activity, reflects differential motor behaviors mediated by distinct cellular and subcellular pools of CB1R. It has been reported that inactivation of CB1R in all cells expressing D1R, including D1-MSNs, impairs the cataleptic response to THC without altering the hypolocomotor response in mice ^45^, and that the pool of CB1R located on D1-MSNs is implicated in THC-induced disruption of motor coordination and catalepsy ^44,46,64^. On the other hand, enhanced locomotor activity in response to amphetamine sensitization, which is reduced in CB1R-deficient mice ^65^, involves CB1R located at striatopallidal D2-MSNs ^46^. This distribution of CB1R activity in D1-MSNs and D2-MSNs is consistent with the classical role of basal ganglia neuronal populations in motor behavior, involving mainly inhibitory and excitatory responses associated with the spiny projection neurons D1-MSNs and D2-MSNs, respectively, although both pathways may be functionally related ^51,53^. Based on this knowledge, we hypothesized that THC-induced hyperlocomotion may involve CB1R in specific striatal neuronal populations. Indeed, our data show that CB1R on membrane terminals of GABAergic D2-MSNs, but not D1-MSNs, are specifically required for THC-induced hyperlocomotion. Our findings are in-line with the well-known action of dopamine on D2R present in the striatum. Indeed, psychostimulants, known to increase dopaminergic tone ^66^, may also target the Akt-GSK3β pathway in the dorsal striatum *via* activation of D2R ^66^. Thus, functional mimicry between CB1R and D2R could ultimately lead to inhibition of the striatopallidal pathway with locomotor disinhibition as the end result. Indeed, a physical interaction between D2R and CB1R has been proposed to occur in striatal cells ^67^. These findings potentially suggest that the two signaling systems might interact to generate specific effects, especially considering that THC can increase the release of dopamine in the basal ganglia. Nevertheless, a role of the CB1R in D1-MSNs in the hyperlocomotor response to a low dose of THC cannot be completely excluded, given that the reduction of CB1R in D1+ cells is not complete in Drd1-CB1R KO mice ^46^.

Interestingly, knowing that phosphorylation at S9 in the N-terminal tail of GSK3β inhibits the substrate association with the substrate binding domain of GSK3β ^31^, we used the viral interference with GSK3β^S9^ phosphorylation in Adora2a-Cre mice to reveal that inhibition of GSK3β activity is a key regulator of THC psychomotor stimulant-induced behavior involving CB1R in the D2 striatopallidal neuronal population. Although ERK1/2^MAPK^ signaling activity has been shown to play a role in striatal motor function in both MSN populations ^68^, no changes in the expression of p-Erk1/2^MAPK^ and total Erk1/2^MAPK^ were observed with increasing striatal THC levels in our experimental conditions. Our results seem to be in line with the work of Hutton and colleagues showing a hyperlocomotor phenotype in mice lacking Erk1/2^MAPK^ in D2-MSNs ^68^, suggesting that an increase in ERK1/2^MAPK^ signaling could be involved in reducing locomotor activity, while as shown here, THC-induced hyperlocomotion seems to depend on CB1R-mediated GSK3β activation in D2-MSNs. In addition, our findings extend the role for GSK3β described in D2-associated stimulation of locomotor activity, as illustrated by the reduction in psychostimulant-induced behavioral activity in mice with GSK3β deletion in D2R-expressing cells ^69^.

### Mechanism of action of PREG and AEF0117 as a therapeutic avenue for THC-induced psychotic-like states

We demonstrated here that PREG and AEF0117 were able to inhibit GSK3β-dependent THC-induced hyperlocomotion (Fig. 5), strongly suggesting a mechanism of action involving β-arrestin1-dependent CB1-mediated GSK3β signaling. The demonstration of a CB1R-mediated β-arrestin 1/Akt/GSK3β-biased modulation involved in the hyperlocomotor effect of THC is highly relevant to the endogenous mechanism of action of CB1R-specific signaling inhibitors (CB1-SSi), namely PREG ^13^ and the first-in-class clinically developed CB1-SSi, AEF0117 ^23^. Our results are consistent with the described role of H8/TM7 in the β-arrestin-mediated, rather than G-protein-mediated, biased signaling state of CB1R ^37,70^. Indeed, PREG and AEF0117 binding to the same allosteric site encompassing TMH1, TMH7 and the intracellular helix H8 of the CB1R were confirmed by their inability to block THC-mediated suppression of cellular respiration in HEK293 cells transfected with human CB1R (hCB1R) when their binding was disrupted by a single point mutation (p.E1.49G) on hCB1R ^13,23^. Although further investigations are needed to demonstrate a direct effect of PREG or AEF0117 on β-arrestin 1-biased CB1R signaling, this unique mechanism of action support a potential beneficial therapeutic potential of AEF0117 ^23^ in psychotic conditions.

In conclusion, our study shows that THC-induced hyperlocomotion at a dose that increases striatal THC levels requires CB1R located at the membrane of striatopallidal neurons and involves inhibition of striatal GSK3β signaling activity through a β-arrestin 1-dependent CB1R signaling pathway. Importantly, this study also identified one of the mechanism of action of CB1-SSi and provides a novel therapeutic target for AEF0117 in cannabinoid-associated psychotic-like disorders.

## Methods

### Animals

Experiments were performed according to protocols approved by the Institutional Animal Care Unit of the INSERM, the Ethics Committee of the University of Bordeaux (CE50, France) and the French Ministry of Higher Education, Research and Innovation (authorization APAFIS#10936; APAFIS#20053) in strict compliance with the French Ministry of Agriculture and Fisheries (authorization B33-12-059) and the European Community Council Directive (2010/63/EU). Every effort was made to minimize animal suffering and to reduce the number of mice used while maintaining reliable statistics.

All experiments were performed on adult male mice, 9-10 weeks old at the beginning of the experiments. Animals were maintained in group housing, and were single-housed in standard plastic rodent cages one week prior to the experiments. Cages and enrichment (cellulose nests and cardboard tunnels) were changed every two weeks. All mice were housed in the animal facility under controlled conditions of temperature (range 20-22°C) and humidity (range 55-60 %) with a constant 12-hour light/dark cycle (light on at 7 am) and *ad libitum* access to food (Standard Rodent Diet A04, SAFE, France) and water.

C57BL/6N mice (weighing 25-30 g) were purchased from Janvier Laboratories (France) and the following mouse lines were generated in our animal facility. β-Arrestin 1 constitutive KO (B6.129X1(Cg)-Arrb1^tm1Jse^/J – Arrb1) mice were purchased from The Jaxon Laboratory (JAX stock #011131). β-Arrestin 1 (-/-) KO mice were obtained by breeding heterozygous male and female β-Arrestin 1 (-/+) mice. CB1-flox mice, generated and maintained as previously described ^46^ were used to generate conditional mouse lines and for the viral deletion experiment. Conditional CB1R mutant mice and their WT littermates were generated by a 3-step breeding scheme using the Cre-loxP system and maintained in our animal facility. To generate the Adora2a-CB1-KO line, CB1-flox mice were crossed with Adora2a-Cre mice (Tg(Adora2a-Cre)KG12633); in Adora2a-Cre mice, provided by MGI (Jackson Laboratory, USA), the Cre recombinase was placed under the control of the adenosine A2A receptor gene (Adora2A) regulatory sequences using BAC transgenesis. Similarly as described above, to generate Drd1-CB1-KO mice, CB1-flox mice were crossed with Drd1-Cre mice (Tg(Drd1-Cre)EY217), in which the Cre recombinase is inserted into the regulatory sequence of the dopamine receptor d1 gene (Drd1a33), provided by MGI (Jackson Laboratory, USA). WT controls for each of these 2 lines were CB1-flox littermates without Cre expression.

### Drugs

THC was purchased as dronabinol resinous oil (ref#THC-1295S-250) or 10 mg/ml solution in 100% ethanol (ref#THC-LOO657-E-1010) from THC Pharm GmbH-The Health Concept (Frankfurt, Germany). The resin was dissolved at 50 mg/ml (w/v) in 100% ethanol. For *in vivo* experiments, ethanol solutions were solubilized in a solution of 4% ethanol, 4% cremophor, 92% saline (0.9% NaCl) and administered to mice by intraperitoneal injection (*i.p.*) in a volume of 10 ml/kg, at a dose of 0.3 mg/kg. For *in vitro* experiments, ethanol solutions were evaporated to dryness, then dissolved in DMSO and used at a concentration of 10 µM for cell treatment. GDC-0084 (5-(6,6-dimethyl-4-morpholino-8,9-dihydro-6H-[1,4]oxazino[4,3-e]-purin-2-yl)pyrimidin-2-amine) (ref#S8163) was purchased from Selleckchem (Cologne, Germany) and was dissolved in a solution of 0.2% methylcellulose, 0.2% Tween80 for oral administration (*p.o.*) via gavage at a volume of 5 ml/kg. D-amphetamine sulfate (Tocris, UK, ref#2813) was kindly provided by Véronique Deroche-Gamonet (Bordeaux, France), dissolved in saline (0,9% NaCl) and injected i.p. at 5 mg/kg in a volume of 10 ml/kg. Haloperidol (Haldol®) was provided by Umberto Spampinato (Bordeaux, France), dissolved in saline (0,9% NaCl) and injected i.p. at 0.1 mg/kg in a volume of 10 ml/kg. Pregnenolone (3β-Hydroxy-5-pregnen-20-one, 5-Pregnen-3β-ol-20-one) (CAS 145-13-1, Sigma-Aldrich) was dissolved in a solution of 1% Tween80, 2% DMSO, 97% saline (0.9% NaCl) to be administered subcutaneously (*s.c.*) in a volume of 10 ml/kg, at a dose of 6 mg/kg. AEF0117 was provided by Aelis Farma (Roowin; batch # L14086-A) and was dissolved in a corn oil solution (Sigma-Aldrich) to be administered *p.o.* by gavage in a volume of 5 ml/kg at a dose of 15 µg/kg.

### Viral Vector and surgery procedures

To generate AAV-DIO-GSK3β and AAV-DIO-GSK3β^S9A^ the coding sequences for HA-tagged GSK3β and the respective S9A mutant (ADDGENE plasmids 14753 and 14754) were cloned in pAAV-CAG-flex plasmid (kindly gifted by Matthias Klugmann, UNSW, Australia) by using standard molecular cloning techniques. The same pAAV-CAG-flex plasmid was used as empty control (AAV-DIO-ctr). AAVs were generated by PEI transfection of HEK293T cells and purified by iodixanol-gradient ultracentrifugation as previously described ^71^. Viral titers were 1.60*10^11^ for AAV-DIO-GSK3β, 6.7*10^11^ for AAV-DIO-GSK3β^S9A^ and 4.51∗1011 for AAV-DIO-ctr, expressed as genomic copies (GC) x ml.

Viral injections of AAV-DIO-GSK3β and AAV-DIO-GSK3β^S9A^ in the dorsal striatum of Adora2A-Cre mice, with respective AAV-DIO-Ctr, were performed with the following coordinates: AP +0,5 L ± 2,0, DV –3.0 as previously described ^72^. Animals were allowed to recover for at least four weeks before the beginning of biochemical (see after) or behavioral experiments. Mice that underwent behavioral experiments were fixed by transcardial perfusion of 4% PFA and their brain were processed for HA immunofluorescence coupled to *in situ* hybridization for D1 or D2 receptor.

### Immunofluorecence and double fluorescent *in-situ* hybridization (FISH) assay

30µm brain slices were cut with a cryostat. Endogenous peroxidases from the cryosections were inactivated with 3% H_2_O_2_ diluted in PBS-DEPC for 30 min at RT. Then, slices were incubated for 20 min at RT in 0.2 mM HCl-DEPC following by an acetylation step of 10 min with 0.1M triethanolamine-HCl (pH 8) and 0,25% acetic anhydride. The riboprobe D1-DIG antisense (1:1000)99 or D2-DIG antisens (1:1000)99 was used to detect mouse Dopamine receptor 1 or 2 mRNA. The riboprobe D1-DIG or D2-DIG sense (1:1000) was used as negative control ^46^. Slices were hybridated overnight at 70 °C with sense or antisense probes. After that, sections were incubated 1 h at RT with anti-DIG POD antibody (1:1500, Roche, ref#11207733910) followed by 10 min incubation with TSA Plus Cyanine 3 (1:100, Akoya biosciences, ref#NEL744001KT). Afterward, sections were permeabilized and blocked for 1 h at RT incubation in blocking solution with 0.3% of triton. Primary antibody rabbit anti-HA (1:1000, Cell Signaling, ref#3724) was incubated over night at 4°C in PBS+0.3% Triton. After some washes, sections were incubated for 2 h at RT in PBS+0.3% Triton with the secondary antibody donkey anti-rabbit AF 488 (1:500, Invitrogen, ref#A21206). Lastly, nuclei were stained with DAPI (1:20000 in PBS). Finally, sections were mounted, dried, coversliped and imaged with a Leica SP5 multi-photon microscope. All DEPC solution had a concentration of 5% v/v DEPC.

### Plasma and brain sampling and THC assay

The brain was quickly removed and brain areas, including the dorsal striatum, were harvested on ice. Blood samples were collected in EDTA-coated tubes. After centrifugation at 500g for 20 min under refrigeration (4 ± 2 °C), the plasma supernatant was removed. All samples were snap frozen and stored at −80 °C until analysis.

THC assay was performed using an isotope dilution method (with internal standard deuterated THC analog, THC-d3) combined with liquid chromatography (LC)-chemical ionization-tandem mass spectrometry (MS/MS) analysis as previously described ^23,59,73^. Mass spectral analyses were performed on a TSQ Quantum Access triple quadrupole instrument (Thermo-Finnigan) equipped with an APCI (atmospheric pressure chemical ionization) source operating in positive ion mode, in conjunction with a Surveyor LC Pump Plus (Supelco C18 Discovery Analytical column) and cooled autosampler. Briefly, samples were homogenized with spiked THC-d3 and the lipid fraction was extracted by using a liquid-liquid extraction with chloroform/methanol/Tris-HCL 50 mM pH 7.5 (2:1:1, v/v) and purified by using a solid-phase extraction (SPE C18; Agilent) with cyclohexane/ethyl acetate (1:1, v/v). The samples were then subjected to LC-APCI-MS/MS in combination with quality control samples to assess between-run and within-run accuracy and precision (CV < 20%) and calibration curve samples (linearity > 0.99) to quantify THC. The mass spectrometer was operated in selected reaction monitoring (SRM) mode to enhance sensitivity, and the concentration of THC was calculated by linear regression of the peak area corresponding to the diagnostic fragment ion (m/z) with the highest intensity.

### Behavioral tests

THC-induced increase in psychomotor stimulation was assessed in a novel environment as previously described ^14^. Briefly, mice were placed in the center of a transparent open field (45 x 35 x 30 cm) with a square-patterned floor (10 cm^2^ each) under controlled light (110 lux). Locomotor activity was assessed by the number of crossed squares and rearings counted during 5 min by the experimenter. A locomotor index including horizontal and vertical locomotion (crosses + rearings) was calculated. As previously described, locomotion was measured 45 minutes after injection of THC or its vehicle combined with 60 min pre-treatment with GDC0084 or its vehicle, with 10 min pre-treatment with PREG or its vehicle ^14^ or with 3 h pre-treatment with AEF0117 or its vehicle ^23^. Independent groups of mice were used for each experimental condition. The locomotor effects of different doses of GDC-0084 were assessed 1 h later in the open field as described above in independent groups of mice. Amphetamine-induced locomotor increase was assessed in the Actimeter apparatus (Imetronic, Marcheprime, France), which consists of a ventilated rack containing 8 individual actimetry cages (30×16×11cm). Each cage was equipped with infrared sensors to detect locomotor activity, and infrared planes to detect rearing. Beam-breaks were recorded and a locomotor index (horizontal and vertical locomotion) was calculated. Amphetamine was administered after one hour of habituation in an actimetry cage, then locomotion was assessed for one hour period (test) as previously described ^46^.

### Cell cultures

One day before the transfection, STH*dh*^Q7/Q7^ cells (CHDI-90000073, Coriell, ref#CH00097) were plated in a 96 well plate (Nunc Thermo Scientific, ref#167008) at 2x104 cells per well in 100µL of complete medium (DMEM - Gibco, ref#61965; 10% FBS not inactivated - Gibco, ref#10270; 1% penicillin/streptomycin - Gibco, ref#15140; 1% G418– Gibco, ref#11811 stock 40 mg/mL). Cells were incubated in a humidified atmosphere at 33°C with 5% CO2. Then STH*dh*^Q7/Q7^ cells were transfected with Lipofectamine 2000 reagent (Invitrogen, ref#11668) in OptiMEM medium (GIBCO, ref#51985) according to the manufacturer’s instructions at a final ratio of 120 µg cDNA / 0.4 µl Lipofectamine per well with *ARRB1* gene (Origen, ref#SC303424) or pcDNA3.1(+). 24h post-transfection, cells were deprived of FBS, penicillin, streptomycin, and G418 and maintained in 100 μl of DMEM (Gibco, ref#61965) per well. After 24 h of deprivation, the culture medium was removed and changed with fresh DMEM (50 µL). The cells were then treated by adding 150 µL of THC solution (40 µM in DMEM, final concentration 10 µM) or its vehicle (DMEM, DMSO 0.2%; final concentration of DMSO, 1.05%). After 30 minutes of incubation at 33°C, the culture medium was removed, and the cells were lysed with 100µL of AlphaLISA lysis buffer (Perkin Elmer, ref#AL003) supplemented with phosphatase and protease inhibitors (1% and 0.5%, v/v respectively; ref#P8340 and ref#P0044; Sigma-Aldrich, USA). The plate was shaken on a plate shaker (350 rpm) for 10 min at room temperature, and samples were stored at −80°C for further alpha analyses ^30^.

### Gene expression analyses

#### RNA extraction

Total RNA was extracted using a standard chloroform/isopropanol protocol and purified by incubation with AmbionTM DNAse I - RNAse free (Thermo Scientific). Purity and concentration of RNA samples were determined from OD260/280 readings using the ND1000 UV spectrophotometer (Fisher Scientific), and RNA integrity was determined by capillary electrophoresis using the RNA 6000 Nano Lab-on-a-Chip kit run on the Bioanalyzer 2100 (Agilent Technologies). cDNA was synthesized from 2 µg of RNA uisng RevertAid Premium Reverse Transcriptase (Fermentas) with a mixture of random primers (Fermentas) and oligo(dT)18 primers (Fermentas). Transcript-specific primers were generated using Primer Express software (Applied Biosystems) based on GenBank sequence information, verified by NCBI BLAST search, and custom synthesized (Eurogentec). Sequences of all primer pairs are listed in Table S1.

#### Real-time quantitative PCR (RT-qPCR)

PCR conditions and LightCycler® 480 SYBR Green I Master mix (Roche Applied Science) were used in a reaction of 10 μl, using transcript-specific primers (0.6 μM) and 2 μl cDNA, corresponding to 4 ng total RNA input. An initial denaturation step at 95 °C for 5 min was followed by 45 cycles of 95°C for 15 sec, 61°C for 15 sec. The PCR data were exported and analyzed using the GEASE (Gene Expression Analysis Software Environment) IT tool developed at the Neurocentre Magendie. The relative gene expression levels of the transcripts were analyzed by the comparative Cq method (2-ΔΔCT) implemented in GEASE, using two reference genes for normalization (Eef1a1, eukaryotic translation elongation factor 1 alpha 1 and Nono, non-POU domain containing octamer binding). Each primer set was tested for the absence of primer-dimer artifacts and multiple products by melting curve analysis and gel electrophoresis. The amplification efficiency of each set was determined by repeated dilution series of a cDNA mixture. Only primers with an efficiency of ∼2 were used.

#### Droplet digital PCR (ddPCR)

The ddPCR mixtures were prepared in a final volume of 20 µl containing the required QX200 ddPCR EvaGreen Supermix (Bio-Rad) with a final concentration of 150 nM of each primer sets and 2 μl cDNA equivalent to 4 ng total RNA input. Each ddPCR assay mixture (20 µl) was loaded into a disposable droplet generator cartridge (Bio-Rad). Then, 70 µl of droplet generation oil (Bio-Rad) was distributed into each of the eight oil wells of the cartridge. The cartridge was then placed in the QX200 droplet generator (Bio-Rad). When droplet generation was complete, the droplets were transferred to a 96-well PCR plate (Dutscher, ref#044701), heat-sealed with foil in a PX1 PCR Plate Sealer (Bio-Rad) and amplified using a Mastercycler Nexus Gradient Thermal Cycler (Eppendorf). Thermal cycling conditions for EvaGreen assays were as follows: 95°C for 5 min, followed by 45 cycles of 95°C for 30 s and 61°C for 1 min, followed by 4°C for 5 min and a final inactivation step at 90°C for 5 min. A no template control and a negative control were included in the assay for each reverse transcription reaction. After thermal cycling the sealed plate was placed in the QX200 Droplet Reader (Bio-Rad) for data acquisition. The resulting data were analyzed using QuantaSoft software (version 1.7; Bio-Rad). For each set of primers, thresholds of fluorescence amplitude were manually set to distinguish between positive and negative droplets. Only the reactions with more than 15,000 valid droplets were used for analysis.

### Brain tissue sampling and protein quantitation

To preserve protein phosphorylation, brain areas were rapidly dissected on ice and placed in dry ice-cold Precellys tubes (Bertin Technologies, Montigny le-Bretonneux, France) stored at - 80°C prior to protein extraction and AlphaLISA assay, as previously described ^30^. Extraction of total proteins from mouse brain structures was performed in AlphaLISA SureFire Ultra lysis buffer supplemented with protease and phosphatase inhibitors (ref#P8340; ref#P0044, Sigma-Aldrich, USA) using the Precellys-24 homogenizer (Bertin Technologies, Montigny le - Bretonneux, France). A two-cycle homogenization protocol (30 s at 5000 rpm) with a 10 s pause between the two cycles using CK14 ceramic beads (ref#03961-1-0032, Bertin Technologies) was used. After homogenization, the samples were centrifuged three times at 10.000 rpm for 10 min at 4 °C, and then the supernatants were quantified. Total protein quantification was performed using a Direct Detect X spectrometer that analyzes the amount of amine bonds (Merck Millipore) by mid-infrared spectroscopy. Protein samples were then stored at −80°C until AlphaLISA analysis.

### AlphaLISA Analysis

A detailed description of AlphaLISA analysis was reported previously ^30^. Total cofilin (ref#*ALSU-TCOF-A10K* and ref#*ALSU-TCOF-A500*), p-Akt1/2/3^T308^ (ref#*ALSU-PAKT-A10K* and ref#*ALSU-PAKT-A500*), p-Akt1/2/3^S473^ (ref#*ALSU-PAKT-B500*), p-GSK3β^S9^ (ref#*ALSU-PGS3B-A500*), p-mTOR^S24/48^ (ref#*ALSU-PMTOR-A500*), p-RPS6^S240/244^ (ref#*ALSU-PS6R-A500*), p-4EBP1^T37/46^ (ref#*ALSU-P4EBP-A500*), p-MEK1^S218/222^ (ref#*ALSU-PMEK-A500*), p-Erk1/2^T202/Y204^ (ref#*ALSU-PERK-A10K*, *ALSU-PERK-A500*), total Erk1/2 (ref#*ALSU-TERK-A10K*, ref#*ALSU-TERK-A500*) levels were quantified using AlphaLISA SureFire Ultra kits according to PerkinElmer’s instructions. Briefly, 10 µl/well of diluted protein samples and 5 μl of acceptor mix (containing both antibodies and Acceptor beads) were added in an OptiPlate-384 white opaque microplate (PerkinElmer, USA). The plate was sealed with a transparent adhesive film and incubated at room temperature for one hour. 5 μl of donor mix (containing Donor beads) was then added into the wells under subdued light (<100 lux). The plate was resealed and incubated at room temperature for one hour. The plate was then read on an Alpha technology-compatible plate reader (EnSpire Alpha plate reader, PerkinElmer). Samples were diluted in lysis buffer to the desired concentration as follows: 0.625 μg / 10 μl for p-Erk1/2^T202/Y204^, total Erk1/2, and p-4EBP1^T37/46^; 1.25 μg / 10 μl for p-GSK3β^S9^, and p-RPS6^S240/244^; 2.5 μg / 10 μl for total cofilin, p-Akt1/2/3^T308^, p-Akt1/2/3^S473^ and p-MEK1^S218/222^, and 5 μg / 10 μl for p-mTOR^S24/48^.

### Data analysis

Analyses were performed using GraphPad Prism software (San Diego, CA). Student’s t-test was used for pairwise comparisons. Multiple group comparisons were examined by one-way or two-way analysis of variance (ANOVA), followed by post hoc analyses using the Dunnett or Tukey multiple comparison test, as appropriate, when significant. The significance level was set at 0.05. For detailed statistical analysis see Figure legends. Behavioral and AlphaLISA data are expressed as % of vehicle.

## Data availability

The datasets generated during and/or analysed during the current study are available from the corresponding author on request.

## Acknowledgments

We thank all the members of the PUMA platforms of the Magendie Neurocentre, in particularly, Delphine Gonzales, Nathalie Aubailly, Elisabeth Huc, Sara Laumond and Julie Tessaire of the animals/genotyping facility for mouse breeding and care; Thierry Leste-Lasserre from the transcriptomics platform for qPCR and ddPCR analyses; Isabelle Matias from the analytical chemistry platform for mass spectrometry analyses; Alexandre Brochard and Benoît Leclerc from the bioinformatics platform. We also thank Yann Rufin from the Biochemistry and Biophysics Facility of the Bordeaux Neurocampus (LABEX BRAIN; ANR-10-LABX-43) as well as the Bordeaux Imaging Center (BIC). We thank Dr Umberto Spampinato, Dr Veronique Deroche-Gamonet and Jean-Francois Fiancette for help in providing reagents. This work was supported by CNRS (to JM.R and M.V.), Inserm (to A.G., V.R.-L., G.M. and L.B.), University of Bordeaux (to A.C., G.D., G.T., P.-L.R.), French State/Agence Nationale de la Recherche (Pregnenoid ANR-22-CE16-0017 to M.V. and L.B., LABEX BRAIN ANR-10-LABX-43 and ANR-23-CE14-0004-03 to G.M., ANR-19-CE14-0039 to L.B., ERA-Net Neuron CanShank ANR-21-NEU2-0001-04 to G.M., CaMeLS ANR-23-CE16-0022-01 to G.M., Hippobese ANR-23-CE14-0004-03 to G.M.), Bordeaux Neurocampus/GPR BRAIN_2030 (Ph.D. Extension Fellowship to P.-L.R.), EU–FP7 (PAINCAGE, HEALTH-603191, to G.M.); the European Research Council (Endofood, ERC–2010–StG–260515; Micabra, ERC-2017-AdG-786467, to G.M.); Fondation pour la Recherche Medicale (DRM20101220445, to G.M., and ARF20140129235, to L.B.); the Human Frontiers Science Program (to G.M.); Region Aquitaine (to G.M.). G.M. and A.B.-G. are recipients of a grant from la Caixa Foundation (agreement LCF/PR/HR23/52430016). The funders had no role in study design, and data analysis and nor in the decision to submit the work for publication.

## Author contributions

M.V., JM.R., PV.P. and G.M. conceived the project; M.V. and JM.R. supervised the project; L.B., A.B.-G., M.Z., JM.R., and M.V designed the experiments; G.T., L.B., PL.R., A.B.-G., V.L., M.Z., V.R.-L., A.C., A.G., G.D., M.M., M.Me., and M.V. performed the experiments; G.T., A.B.-G., M.Z., G.D., L.B., JM.R., and M.V. analyzed and interpreted the data; M.V. wrote the manuscript; M.V., JM.R., L.B., G.M. and PV.P. revised the manuscript. All authors approved the manuscript.

