## Supplemental Figure S1, Figure S2, Figure S3, Table S1, Figure S4, Figure S5, for "Pregnenolone and AEF0117 block cannabinoid-induced hyperlocomotion through GSK3β signaling at striatopallidal neurons"

**Tomaselli, Bellocchio et al. - Supplementary**

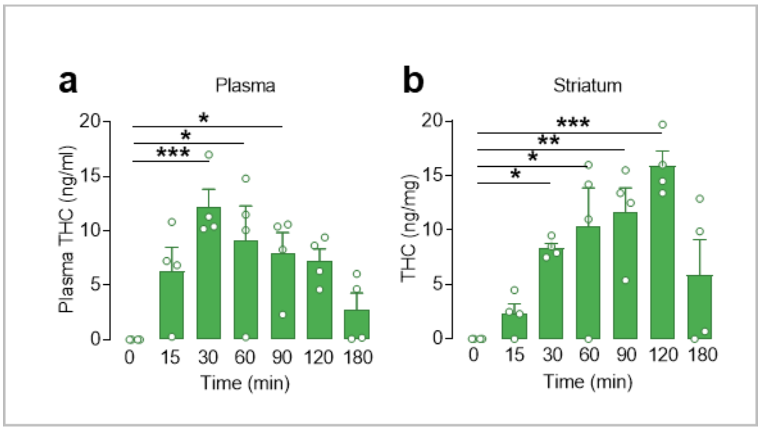


**Fig. S1. THC injection increased THC levels in the plasma and striatum of mice over time.**

**a**. Plasma THC levels peaked 30 min after i.p. injection of THC at 0.3 mg/kg and remained elevated for 90 min. One-way ANOVA, F(6, 21)=4.69; P<0.01; Dunnett's multiple comparison test, T0 vs. T30 min, ***p<0.001, T0 vs. T60 min, *p<0.02 ; T0 vs. T90 min, *p<0.05 ; T0 vs. T120 min, p=0.056 ; T0 vs. T180 min, p=0.79. **b**. Striatal THC levels increased from 30 to 120 min after i.p. injection of THC at 0.3 mg/kg. One-way ANOVA, F(6, 21)=6.81; P<0.001; Dunnett's multiple comparison test, T0 vs. T30 min, *p<0.05 ; T0 vs. T60 min, *p<0.02 ; T0 vs. T90 min, **p<0.01 ; T0 vs. T120 min, ***p<0.001 ; T0 vs. T180 min, p=0.24. Data are mean ± SEM of single values.

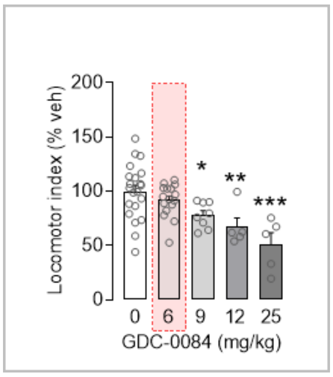


**Fig. S2. GDC-0084 induced a dose-effect decrease in locomotion in C57Bl/6N mice**. Locomotion is measured in an open-field 1 h after oral administration of GDC-0084 at 6, 9, 12 or 25 mg/kg or vehicle (0 mg/kg). The locomotor index including horizontal and vertical locomotion (crosses + rearings) is expressed as a percentage of the vehicle. One-way ANOVA, F(4,50)=7.6, ***p<0.001; Dunnett's comparison multiple test, GDC-004 at 6 mg/kg: p=0.64, 9 mg/kg: *p<0.05, 12 mg/kg: **p<0.01, or 25mg/kg: ***p<0.001 vs. vehicle (0 mg/kg). The 6 mg/kg dose was ineffective on locomotion and was therefore chosen to study the effect of GDC0084 on THC-induced locomotion (see Fig. 2).Data are mean ± SEM of single values.

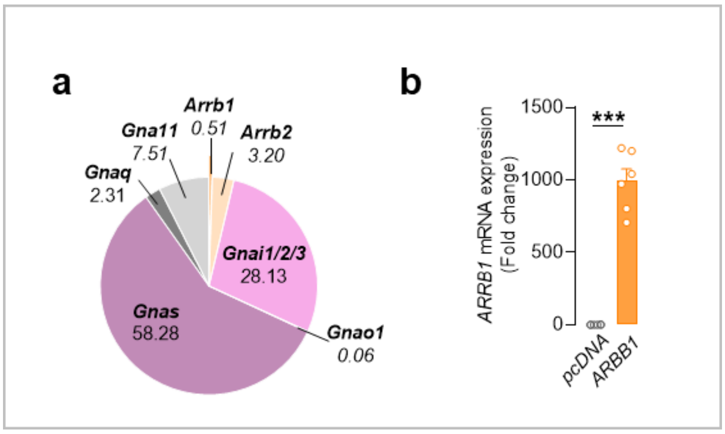


**Fig. S3. PCR analysis of β-arrestin and G-protein encoding genes in STH*dh*$^{Q7/Q7}$ cells.**

**a.** Relative percentage of mRNA expression of β-arrestin (*Arrb1* and *Arrb2*) and G-protein (*Gnai1/2/3*, Gnao, *Gnaq*, *Gna11 and Gnas*) encoding genes measured by droplet digital PCR (ddPCR) in STH*dh*$^{Q7/Q7}$ cells. **b**. *ARRB1* mRNA expression measured by real-time quantitative PCR (RT-qPCR) in STH*dh*$^{Q7/Q7}$ cells after 48 h post-transfection with human *ARRB1* encoding gene or with empty pcDNA. Two-tailed unpaired t-test, *ARRB1* vs pcDNA, t=12, df=10, ***p<0.001. Data are mean ± SEM of single values.

**Table. S1. List of forward and reverse primers used for PCR analysis.**

| Gene name | GenBank ID° | Forward (5'-3') | Reverse (5'-3') |
|---|---|---|---|
| **Arrb1** | NM_177231 | CCCCATGTGTGAAGGGCTAGTA | TCACCCTCATATCCCTTTCAGTAGT |
| **Arrb2** | BC016642 | CCGCTATGGCCGAGAAGAC | CTGGTAGGTGGCGATGAACA |
| **Gnai1** | NM_010305 | TCGTGCCATTGAAACAAAATCA | TGGGAAGGCGAGTCAGCTT |
| **Gnai2** | NM_008138 | GCCGGGAGTAGCCATGGTA | CCCCAGAACAAGACAACGGTTA |
| **Gnai3** | NM_010306 | GCTGTGGCTCGGATCCATT | CAGTTCTGACCATCAACCTTCTTTC |
| **Gnao1A** | NM_010308 | AAGCCCACGGGTCTTTGTAA | AGGCTGGCAGTGTCAGGTGT |
| **Gnao1B** | NM_001113384 | TTCTGGGCCTCCCGGTAA | GAGGTGGAGCCTCAGGACTG |
| **Gnaq** | NM_008139 | CTTCTTAGGAGCCTGCGTATATTGT | GTTCCAGGTGTTCTTTAAATATCGC |
| **Gna11** | NM_010301 | CCACCTTCCCACACTAGGCTC | TGTAGTGTATGGATGGCACAGAT |
| **Gnas** | NM_201618 | GTGACCCGGGCCAAGTACT | TGGCGCCCATCTCCACTA |
| **Reference genes** | | | |
| **Eef1a1** | NM_025380 | TGAAAATCAGTGGCTCAACTTTAAA | ACACGAAAAAAGTAAACAGCCAGA |
| **Nono** | NM_023144 | CTGTCTGGTGCATTCCTGAACTAT | AGCTCTGAGTTCATTTTCCCATG |

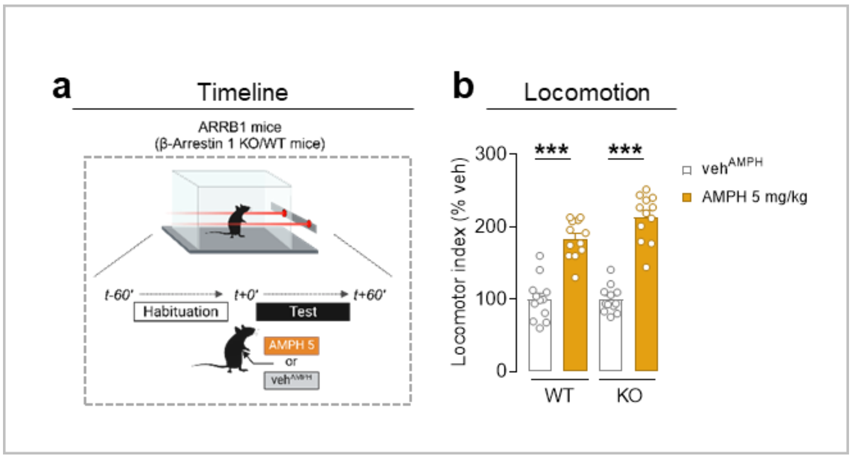


**Fig. S4. Amphetamine-induced hyperlocomotion in β-arrestin 1 knock-out (KO) mice.**

**a**. Timeline of the locomotor activity procedure in β-arrestin 1 knock-out (KO) mice and wild-type (WT) littermates in the actimetry cages. Amphetamine (5 mg/kg) (AMPH 5) or vehicle (veh$^{AMPH}$) was administered i.p. after 1 h of habituation in the actimetry cages, then locomotion was assessed for 1 h (test period). **b**. Effect of amphetamine on locomotor activity in WT and KO mice during the test period. The locomotor index, including horizontal and vertical locomotion, is expressed as a percentage of veh$^{AMPH}$. Two-way ANOVA, treatment effect, $F(1,44)=159$, $p<0.001$; Tukey's multiple comparison test; AMPH vs. veh$^{AMPH}$, in WT and KO mice, ***$p<0.001$. Data are mean ± SEM of single values.

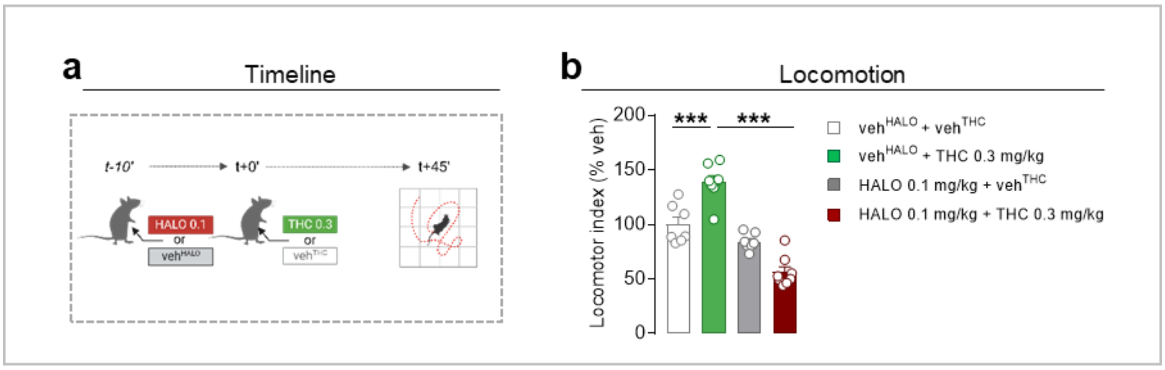


**Fig. S5. Haloperidol blocks the effects of THC on locomotion.**

**a**. Timeline of locomotor activity measurement in an open-field 45 min after i.p. injection of THC (0.3 mg/kg) or vehicle (veh$^{THC}$) in mice with a 10 min pre-treatment with i.p. injection of haloperidol (HALO) at 0.1 mg/kg or vehicle (veh$^{HALO}$). **b**. Effect of haloperidol on locomotion in mice treated with THC or vehicle (veh$^{THC}$). The locomotor index including horizontal and vertical locomotion (crosses + rearings) is expressed as a percentage of the vehicle. Two-way ANOVA, treatment interaction, $F(1,26)=39.3$, $p<0.001$; Tukey's multiple comparison test, THC vs. veh$^{THC}$ in mice treated with veh$^{HALO}$, ***$p<0.001$. HALO effect vs. veh$^{HALO}$ in THC-treated mice, ***$p<0.001$. HALO effect vs. veh$^{HALO}$ in veh$^{THC}$-treated mice, $p=0.2$. Data are mean ± SEM of single values.
